# International Mouse Phenotyping Consortium: Investigating gene function and providing insights into human disease

**DOI:** 10.1101/2025.10.09.681205

**Authors:** Robert Wilson, Tuğba Bülbül Ataç, Tsz Kwan Cheng, Anthony Frost, Osman Güneş, Marina Kan, Piia Keskivali-Bond, Federico López Gómez, James McLaughlin, Jakub Mucha, Tawanda Munava, Carla Oliveira, Diego Pava, Jose Francisco Peña Estrada, Ewan Selkirk, Bora Vardal, Sara Wells, Pilar Cacheiro, Damian Smedley, Helen Parkinson

## Abstract

The International Mouse Phenotyping Consortium (IMPC; https://www.mousephenotype.org/) web portal contains phenotype data for mouse protein-coding genes derived from analysis of data obtained in a systematic and high-throughput fashion from knock-out lines produced by IMPC. The project has produced >1,400 mouse models of human disease that recapitulate phenotypes observed in patients. Over 8000 papers rely on data or reagents generated by IMPC, demonstrating the impact of the project on the research and clinical communities, and IMPC data is incorporated into other resources, such as MGI, Open Targets and UniProt. Data release (DR23.0, 2025) contains > 100 million data points from 9,277 genes and identified 113,803 significant phenotypes. To manage efficient access to this quantity of high dimensional data the IMPC web portal has been rebuilt using a cloud native architecture. The modern user interface retains the look and feel of the original portal with improvements identified through a usability study. New data visualisation and training materials for large scale data access through the API have also been developed to make the resource easier to use.

**Graphical abstract:** 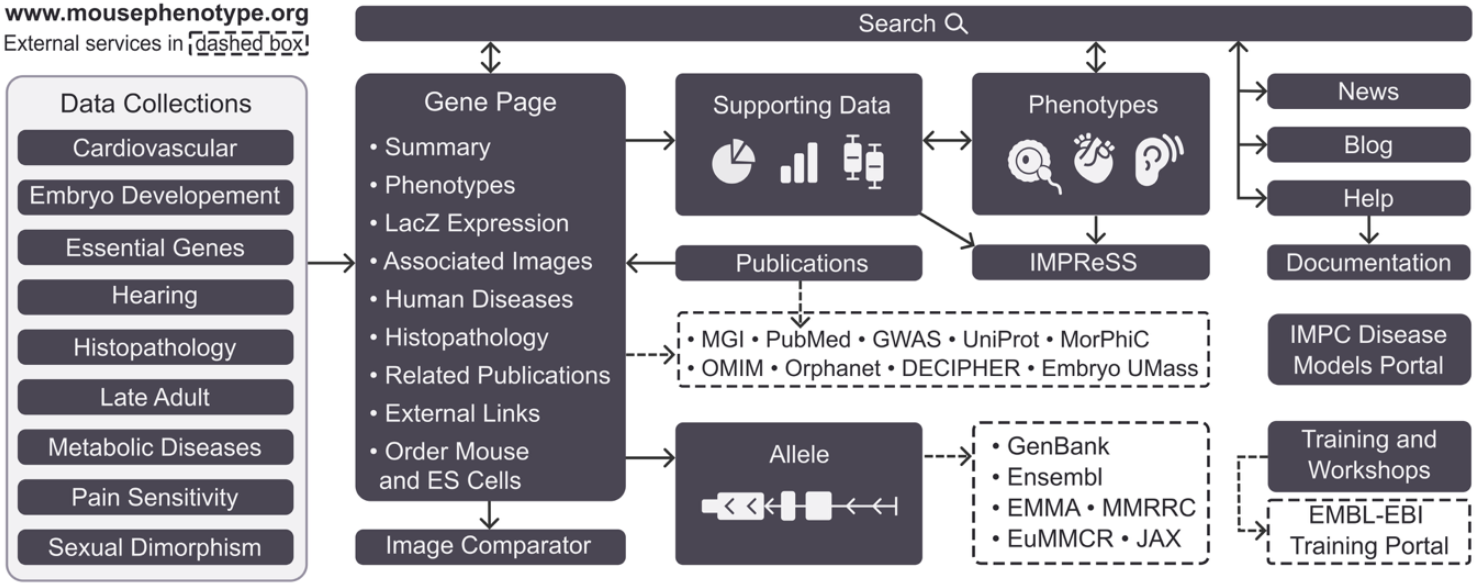

## Introduction

The International Mouse Phenotyping Consortium, (IMPC) is a global effort established in 2011 that involves 21 institutes in 15 countries who are collaborating to generate a comprehensive catalogue of mammalian gene function by knocking out and systematically phenotyping every protein-coding gene in the mouse on an inbred C57BL/6N genetic background (1,2,3). The project provides a comprehensive catalogue of mammalian gene function and improves understanding of human disease via >1400 models of human disease genes (Disease Models Section) which recapitulate the phenotypes seen in human patients. IMPC data has been used to identify >100 validated rare disease disease-gene associations and (4) opened new avenues to explore for the therapeutic treatment of diseases (5,6,7,8). IMPC phenotyping protocols are standardised and harmonised across the project (3) and the phenotyping assays provide reproducible, sex and life stage phenotypes for 16 categories, including anatomy, clinical chemistry and ageing. Genes studied by IMPC are prioritised for targeting by using an unbiased algorithmic strategy, most recently targeting under studied genes (9,10,11); Gene essentiality is the primary factor influencing the ability to generate a null allele (12) and analysis of IMPC viability data and comparison of IMPC data with e.g. cell derived datasets has, provided a framework to investigate the spectrum of intolerance to loss-of-function variation by binning genes into essentiality categories: cellular lethal, developmentally lethal, subviable, viable with phenotype, and viable with no phenotype (13). These gene essentiality categories align with human disease gene classifications, and refinement of gene essentiality analysis (14) has helped prioritise predicted pathogenic variants in genes not currently linked to Mendelian conditions, enhancing the approach that has already been shown to improve the genetic diagnosis of human diseases (15).

The IMPC portal is used by biomedical researchers, common and rare disease researchers, data scientists, informatics users and resource developers (3, 16). Information about the IMPC project is disseminated under the CC-BY 4.0 licence through the web portal, API, FTP data downloads and a knowledge graph (3). These resources provide the scientific community with free and unrestricted access to the primary and secondary data, the genotype-phenotype annotations made by IMPC, disease associations, the standard operating protocols used to perform the broad based phenotyping of the animals, and links to stock centres where the mice and reagents generated by IMPC can be obtained. The latest release (DR23.0, 2025), comprises >100 million experimental observations for 9,277 genes with 113,803 significant phenotype calls (51,190 for embryonic stages, 57,918 for early adults < 16wk, and 4,695 for mid/late adults) (17), assayed in homozygotes, hemizygotes and heterozygotes animals from 9,994 independently established lines. Statistics presented in subsequent sections derive from DR23.0 unless otherwise stated. In this article we focus on, new approaches to identification of disease models, web portal improvements addressing user feedback, portal sustainability and data visualisation, improved training tools and materials for scalable data access and improvements to data capture and standardisation that have enhanced the quality of the data. These changes to the portal have made it more sustainable, easily extensible and better placed to showcase the large quantity of high dimensional data that IMPC produces.

### Improved data acquisition

Data acquisition and QC related processes and reporting have been improved with services moving under a unified interface with harmonised API calls to track data through the different processes. The interface allows easier generation of new reports and modification of existing ones that are automatically updated for the data generating centres. This strategy has been utilised for QC reporting and alongside automation of some QC checks and standardisation of issue titles and definitions allows faster resolution of QC issues. Updates have also been made to the data completeness process and reporting making it easier to ensure all data has been submitted. These processes and reports ensure faster turnaround of the data for the IMPC portal.

### Disease Models

The IMPC viability data and the gene essentiality analysis have revealed associations between essentiality categories and specific types of disorders (13,18), ultimately leading to the identification of novel genes associated with neurodevelopmental disorders (19,20). In addition to capturing Mendelian disease associations from OMIM (21) and Orphanet (22) for the corresponding human orthologous genes, the automated identification of mouse models of human disease is facilitated by the PhenoDigm algorithm (23). This algorithm calculates a similarity score for all pairwise combinations of mouse knockouts and human single gene disorders that reflects the ability of the mouse model to mimic the phenotypes observed in humans. In DR23 there were 2,938 mouse models for human one-to-one orthologs associated with known Mendelian diseases. The human disease pipeline requires association of MP annotations with the mouse knockouts and HPO terms for diseases, resulting in analysis of 2,581 genes by the PhenoDigm algorithm and the identification of 1,435 mouse models that recapitulate some of the human disease phenotypes. Several factors influence the ability of mouse models to mimic the clinical features of human disease. These include the absence of assays for certain phenotypes, the pleiotropy and severity of the disease, differences in physiology and gene regulation between the two organisms, as well as variations in viability and zygosity of the disease-matched mouse knockout (see 4). A summary of the similarity scores to human disorders associated by orthology, as well as potential novel disease associations based on phenotypic similarity, is displayed on the portal for each gene (Figure 1), and can also be accessed and downloaded in bulk through the IMPC disease models portal (https://diseasemodels.research.its.qmul.ac.uk/) (4).

**Figure 1.**
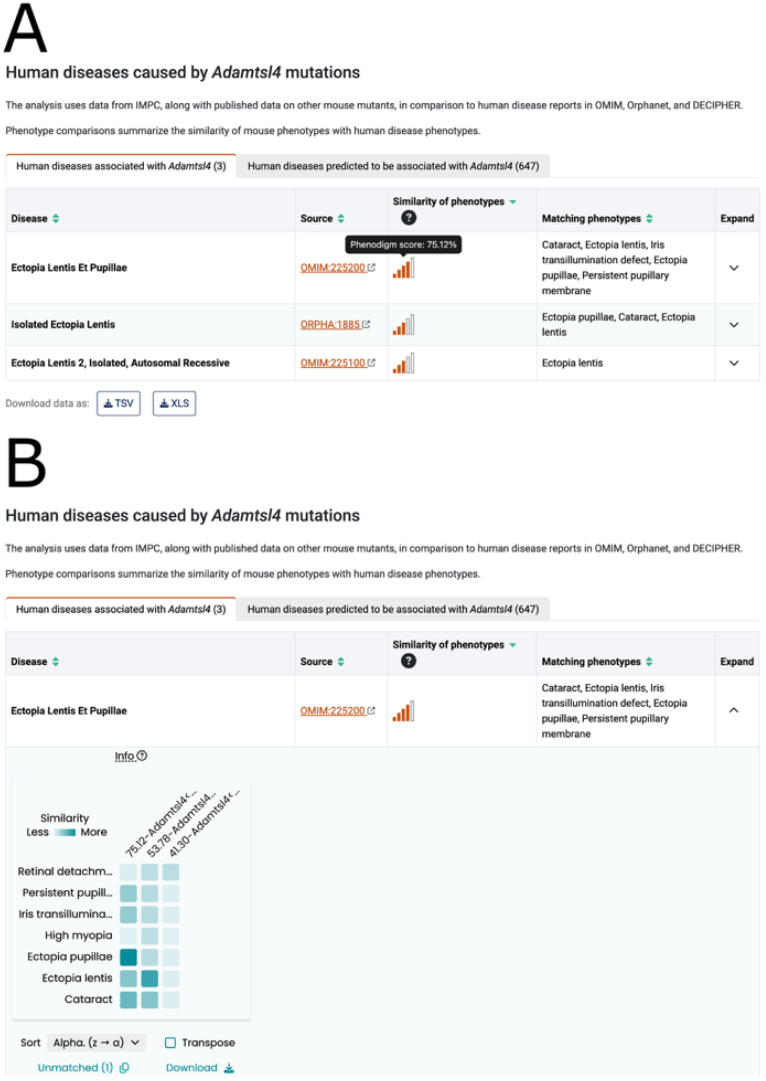
Example human disease section for *Adamtsl4*. (**A**) The left tab contains human diseases associated by orthology as curated by OMIM, Orphanet, and/or Decipher. The disease name, identifier, PhenoDigm score, and human phenotypes matching the mouse model’s phenotype are displayed on the table. The right tab contains human diseases predicted to be associated with the gene through phenotypic similarity. (**B**) The Phenogrid is a collaboration between the Monarch Initiative and the IMPC. It displays the phenotypic similarity of individual phenotypes between the human disease (y-axis) and the matched mouse models (x-axis) as a grid. In this example, the match with the highest similarity score for the autosomal recessive human disease Ectopia lentis et pupillae (OMIM:225200) and its associated gene, *ADAMTSL4*, was the IMPC orthologous late adult mouse model [*Adamtsl4*<*em1(IMPC)J*> *hom late*] with a score of 75.12%. Darker boxes indicate a higher similarity between human and mouse phenotypes, for example, *Ectopia pupillae* (HP:0009918) has a perfect match with *abnormal placement of pupils* (MP:0006241). Further information on PhenoDigm scores available on the help and data analysis pages:https://www.mousephenotype.org/help/data-visualization/gene-pages/disease-models/; https://www.mousephenotype.org/help/data-analysis/disease-associations/

### Web portal Improvements

#### Infrastructure

The portal has been rebuilt as a cloud native application designed to address the current scale of the data/access needs and offer improvements to portal usability and extensibility. To improve page load times the portal uses the component model in which each section of the page loads data from a separate microservice in parallel. We also reviewed the structure of the data to optimise the information for each section so the data could be fetched in a single request. Where a significant amount of data was required, for example for the disease section of the gene page, we fetched a small amount of data initially and loaded the rest after the page had completed loading. These strategies significantly improved the performance of the pages for users.

### Design and Navigation

The portal design was conserved to retain the look and feel for users. User experience (UX) testing with 13 existing and new users and a usability review (see https://www.nngroup.com/articles/ten-usability-heuristics/) provided user-led feedback and retesting with users after revision improved our design. Consistency of design for links and breadcrumbs was reviewed, and the visibility of buttons, labels, tables and forms was increased by improving colour contrast, which is a simple but effective way to enhance the communication of information. Improved visibility of gene-phenotype p-values for statistical data and new filters and sort functions were added for data tables which improved the user experience especially for large tables. Site navigation was also improved as the IMPC content has grown over time and become complex. New features were evaluated using moderated usability testing with a subset of the first UX group to determine if the site was sufficiently improved.

### Data Presentation and Navigation

IMPC’s data is large and complex, therefore the summarisation of phenotypes found for each mouse gene was improved with increased visibility for links to deeper information elsewhere on the page. For example, users expect to see a link to physiological system(s) when significant phenotypes are present (Figure 2A). The ability to order materials from IMPC’s associated stock centres (MMRRC (24) and EMMA (25)) is a requirement for users, therefore this user journey has been redesigned, with a newly located and named button ‘Order Mice’ on gene query results page and a new ‘View Allele Products’ button at the top of each gene page. The increased visibility of these links along with links to other sections on the gene page offers users rapid navigation and easier access to data of interest.

**Figure 2.**
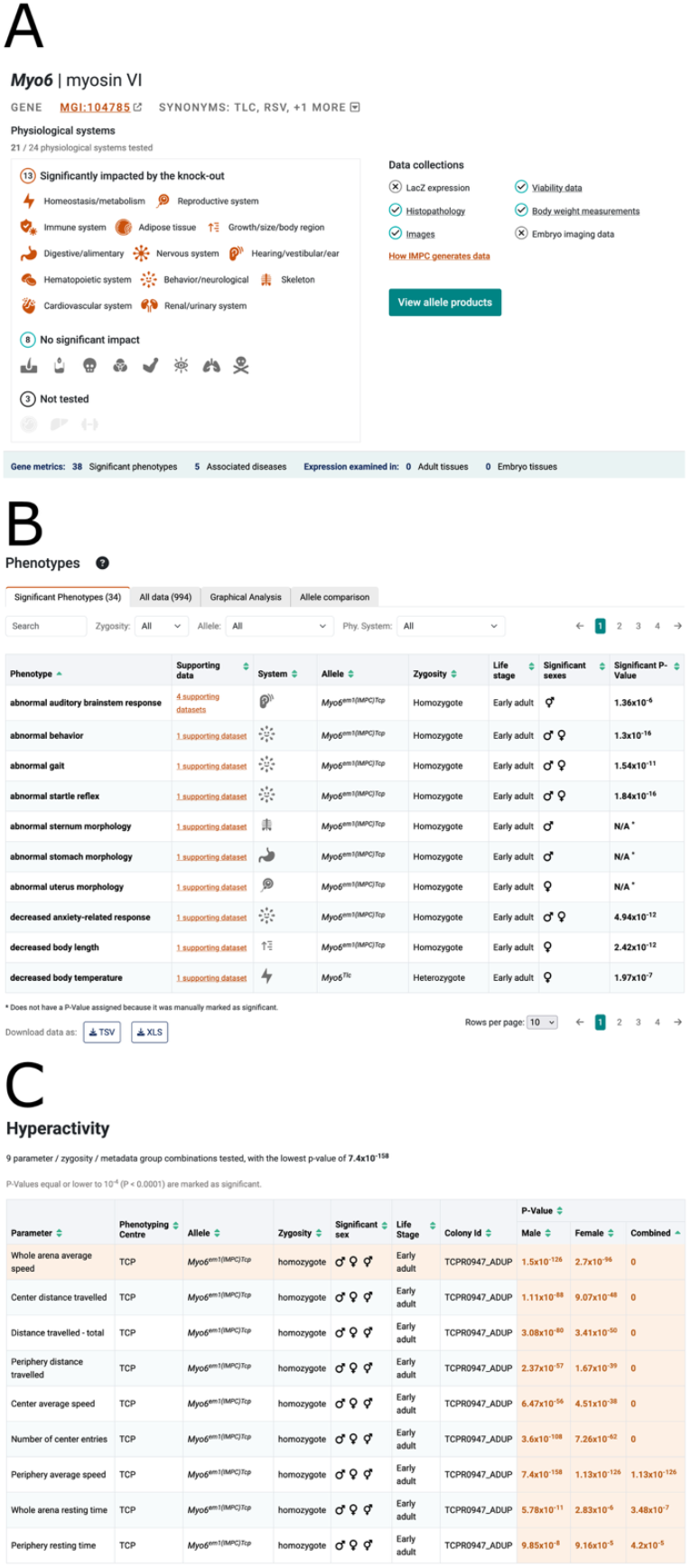
Improvements to the gene report pages. (**A**) The summary section for the *Myo6* gene report page showing the physiological systems impacted by the knock-out, which link to the phenotypes section of the page to show the data in for the system. Links to data collections for the gene, such as body-weight and viability data, also improve navigation and access to allele products is more prominent. (**B**) The phenotypes section contains a number of tabs to explore IMPC data. The significant phenotypes tab contains a prominent link to the data supporting the phenotype and filters for zygosity, allele and physiological system make it easier to identify phenotypes of interest. (**C**) The table summarising all experimental observations and p-values at the top of the supporting data page. In some cases, such as hyperactivity, multiple experimental observations support the phenotype call. Clicking on a row in the table shows the data for the experimental observation below the summary table. Significant p-values are shown in orange text and the p-value threshold is stated above the table.

For the phenotype section we made several key changes to simplify navigation to the supporting data that were guided by usability testing. We added an explicit link to the data supporting the phenotype association for the specific gene, which was previously accessed by clicking on the row, and placed this link next to the phenotype term to increase its visibility. We also added a filtering system to this section to enable selection of phenotypes by physiological system in response to feedback during testing (see Figure 2B). These changes make it easier for users to find the supporting data behind the phenotype calls and identify phenotypes of interest.

On the gene page we report the most significant p-value identified for the phenotypes. IMPC, however, evaluates sexual dimorphism (26), and there is not always a simple one-to-one association between a phenotype and an experimental observation. For example, a hyperactivity phenotype can be assigned based on any one of a number of different open field test measurements that assess anxiety and exploratory behaviours, such as centre average speed and periphery distance travelled. To clearly communicate the data supporting the phenotype assignment we now indicate the number of observations in the supporting datasets link, and summarise all experimental observations and p-values in a table at the top of the supporting data page (Figure 2C). Based on user feedback we clarified the p-value threshold that IMPC uses to assign a phenotype, and significant p-values are highlighted in the summary table to better communicate them to users. Selection of a row in the summary table makes it active, updating the section below to display the charts presenting the results for that particular experimental measurement. This reorganization of the supporting data page keeps the page compact and provides users with an immediate overview of how the experimental data supports the phenotype call.

### Improved graphical analysis tools

Experimental observations are plotted in relation to the magnitude of the p-value (Figure 3 A). Each dot is coloured according to the physiological system, and the p-value threshold used to make a phenotype call is shown as a dotted line. Hovering over a dot provides more data, including: the number of mutants, zygosity and effect size. Zooming in and out and scrolling allow intuitive and dynamic exploration of the data.

**Figure 3.**
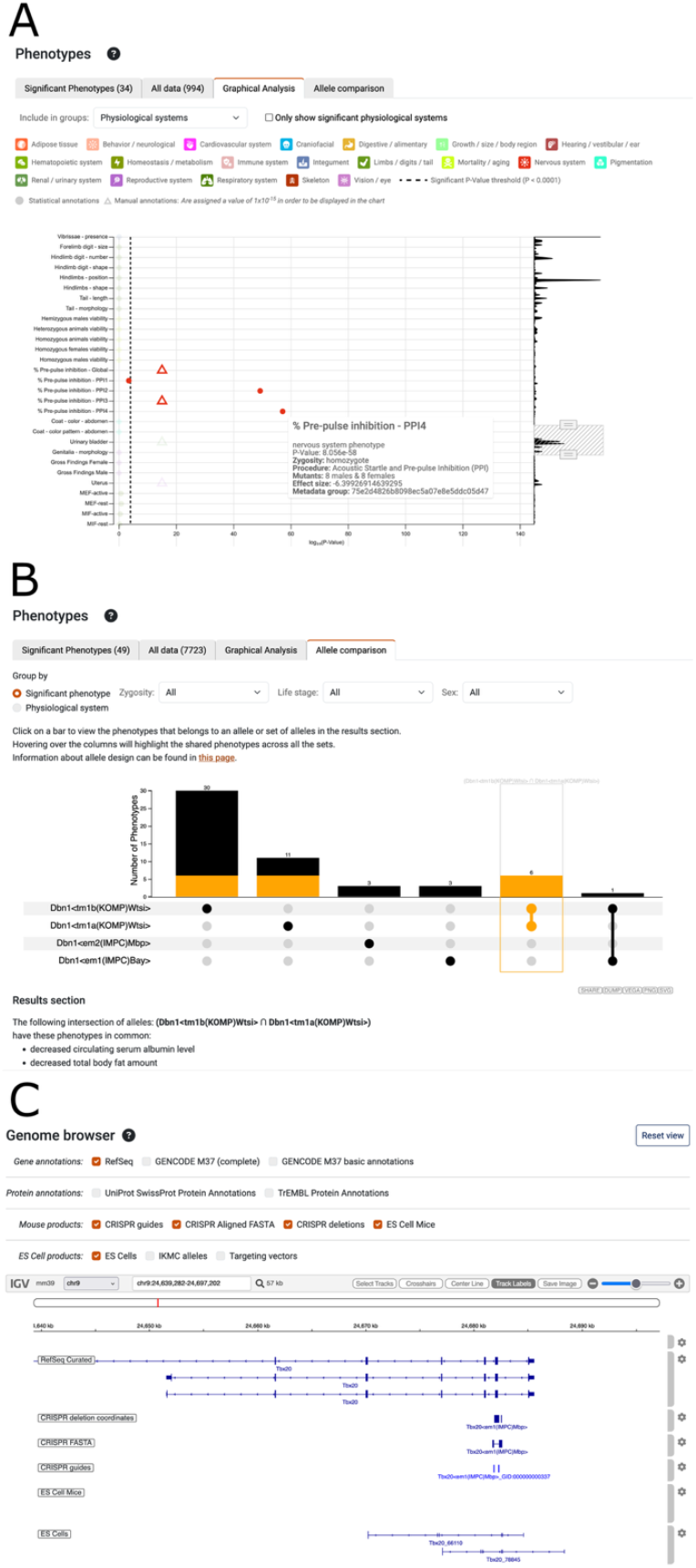
New and improved visualisation. (**A**) The graphical analysis tab of the phenotypes section displays the magnitude of the p-values for experimental observations in relation to the p-value threshold used to assign phenotypes. On the right a resizable sliding window allows changing the region viewed and the zoom level. An information panel, as shown for pre-pulse inhibition - PPI4, is displayed when hovering over a data point. Manual annotations are shown as triangles and assigned an arbitrary value of 1×10^−15^ to include them on the chart. (**B**) When more than one allele has been examined the Allele comparison tab enables phenotypes to be compared for different combinations of alleles, represented as the set of connected dots below the histogram. Selecting a bar in the histogram, as shown for *Dbn1*<*tm1b(KOMP)Wtsi*> and *Dbn1*<*tm1a(KOMP)Wtsi*>, displays the phenotypes for that set below the graph. Filters for zygosity, life stage and sex enable different slices of the data to be selected, and the phenotypes can be summarised to compare across physiological systems. (**C**) The genome browser reveals how IMPC alleles impact gene models. RefSeq gene models are shown by default, but optionally GENCODE, UniProt SwissProt and TrEMBL annotations can be displayed. For IMPC CRISPR alleles, tracks show the location of the CRISPR guides, alignment of the FASTA sequence and the calculated deletion. For ES Cell alleles the IKMC allele track from the UCSC genome browser was migrated from the mm9 to the GRCm39 assembly. By default, only the ES Cell derived products available to order through the IMPC website are shown.

### Phenotype comparisons

Some IMPC genes have multiple alleles, and these can be associated with different phenotypes. Therefore, we developed a comparison tool that enables users to see shared phenotypes (Figure 3B). UpSet visualisation was selected due to complexities of the data (34). A histogram indicating the number of phenotypes is shown for each allele, indicated by a solid dot in the table, and for different combinations of alleles, shown as a line intersecting solid dots. Filters enable refinement of the data according to the zygosity of the alleles, the life stage or the sex of the animals. To make it more flexible users can compare the data by the physiological system impacted by the phenotypes as well as by the specific phenotype terms associated with the alleles (Figure 3B).

### Visualisation of ES and CRISPR derived alleles

IKMC ES cell derived alleles, used to generate IMPC mouse lines, are currently visible via a track on the UCSC browser (https://www.genome.ucsc.edu/cgi-bin/hgTrackUi?hgsid=2971680668_bMXqX3P9MNhX4mrOACbIA6FXbPAy&db=mm9&g=ikmc), however, the NCBI37/mm9 assembly is outdated and none of the IMPC CRISPR derived alleles are represented, meaning users could not assess the impact of deletions vs. current genome annotations. Therefore the data for the ES cell derived alleles were migrated to GRCmm39 using LiftOver and CRISPR derived alleles guides were aligned to the genome using BLAT (27) and BedTools (28) and deletion coordinates were derived using BEDOPs (29). The tracks are displayed on allele pages using Integrative Genomics Viewer (30) with RefSeq (31), Gencode (32) and UniProt (33) annotations (Figure 3C). A track hub is also available https://ftp.ebi.ac.uk/pub/databases/impc/other/impcTrackHub/hub.txt for use in UCSC and Ensembl genome browsers. These resources enable users to evaluate the exons deleted and whether the allele is likely to represent a null given the current gene models.

### Improved programmatic data access

In conjunction with rebuilding the IMPC website we have improved and simplified programmatic access to the IMPC data. We have focused on helping users to integrate IMPC data into their workflows by reducing steps required between data access and analysis and widening access for non technical users. IMPC data is available from an Apache Solr API service. To make it easier for users to access IMPC data programmatically (without knowledge of SOLR) a Python library (https://github.com/mpi2/impc-api) now wraps the API and a training course “Accessing Mouse Phenotypes and Disease Associations with the IMPC Solr API” (https://www.ebi.ac.uk/training/online/courses/impc-solr-api/#vf-tabssection--contents) explains how to use the library and outlines common query patterns and best practice.

### Data Archiving

IMPC has gathered a unique collection of 852K images collected from 28 different procedures during the phenotyping process. The set of 465K Xray images are now archived in the BioImage Archive (BIA) (https://www.ebi.ac.uk/biostudies/studies/S-BIAD2244), and we are in the process of depositing the remaining images. The BIA is a permanent repository for image data (35) ensuring they are available for the wider image analysis user community to use as a reference dataset for AI and to ensure long term availability of complex and large image datasets.

## Discussion

The quantity of high quality IMPC data that has passed a rigorous QC process, see (3) and the improved data acquisition section, has grown by 15 million data points since 2023 to just over 100 million data points and is anticipated to continue to grow at a similar rate. Modernisation of the portal means the resource is now in a much better position to be able to manage this quantity of high dimensional data while retaining a performant user interface. The new infrastructure has made the resource more sustainable, docker images enable deployment of the resource in different cloud environments and the microservice architecture makes it easier to add new features. The modern user interface keeps all the functionality and look and feel of the original site, while improving the graphical analysis tool and adding new features, such as the genome browser and the allele comparison tool. A usability study to evaluate the interface improved the layout of the summary section, resulting in an effective path to finding allele information, a key function of the site, and helped improve the navigation between phenotypes and supporting data. Testing also refined the layout and communication of information producing a web portal that is easier to use. Accompanying the revision of the website the documentation and online training courses were updated and improved, notably for programmatic access of data through the API for which a Python library was created to make it easier to access the data.

IMPC tracks the impact of the project through papers published involving reagents generated by the project, currently > 8200 have been curated. This helps understand communities using IMPC data, which potentially could help foster collaborations. For example, a tool to quantify the asymmetry in embryos and assess the contribution of genes to abnormal asymmetry has recently been developed based on IMPC 3D image data that could help understand developmental disorders and neurological disorders including schizophrenia, autism spectrum disorder, and Down’s syndrome (36). A summary of the impact of the publications identified though the IMPC tracking system along with additional examples involving biomarker discovery, investigation of gene therapy interventions and drug efficacy among others has recently been published (8).

While the data for each IMPC data release is available from the EBI FTP site, the deposition of IMPC image data into the BioImage Archive provides an additional way to disseminate IMPC data and will promote further reuse of the data by a wider audience, including image machine learning specialists. IMPC data is also incorporated into other resources, such as MGI, Open Targets and UniProt increasing access to the data for a wider user community. For example, IMPC phenotype data in Open Targets has been used as part of the validation strategy to prioritise candidate disease genes based on tissue specific protein-protein associations (37). The IMPC track hub, which can be loaded into either UCSC or ENSEMBL genome browsers, offers users of those resources the ability to view IMPC alleles in the context of other genomic features or their own experimental data.

The knowledge graph, that includes IMPC and GWAS data, could be used to add new functionality to the IMPC web portal. While the current section on human disease associations and computational identification of disease models is centred around Mendelian, single-gene disorders, ongoing efforts aim to map mouse phenotypes to EFO-encoded GWAS traits and integrate this information into the portal. For example, data could be extracted from the knowledge graph to power a disease based search based on links to GWAS data and human orthologs of mouse genes. The IMPC knowledge graph is also being used to add IMPC mouse phenotype data to other knowledge graphs, such as the CFDE Data Distillery Knowledge Graph and UniProt ProKN knowledge graph. To further enhance the distribution and reuse of IMPC data we will ensure IMPC data is presented effectively to large language models (LLMs) via a Model Context Protocol (MCP) server and explore how we can integrate the knowledge graph with LLMs. One potential application of a LLM for the website would be to build an interactive help desk, as has been done for other resources such as PRIDE (38), to provide users with a fast response to questions and an alternative way to access the web site documentation. The value of engaging your user community to help with the design of the website should not be underestimated, and it is important to ensure existing features work well and are valued by users before considering adding new features.

## Data availability

- IMPC Web Portal: https://www.mousephenotype.org/
- IMPC FTP data releases: http://ftp.ebi.ac.uk/pub/databases/impc/
- BioImage Archive data: https://www.ebi.ac.uk/biostudies/studies/S-BIAD2244
- IMPC Knowledge Graph: https://ftp.ebi.ac.uk/pub/databases/spot/kg/impc_kg_neo4j.tgz
- IMPC Track Hub: https://ftp.ebi.ac.uk/pub/databases/impc/other/impcTrackHub/hub.txt
- IMPC disease models portal: https://diseasemodels.research.its.qmul.ac.uk/

## Supplementary data

None

## Acknowledgements

We are extremely grateful to the individuals who gave up their time to help us with the usability study and thank the members of the IMPC and all users of our services. We thank Tudor Groza for initiating the cloud native project and Jonny Lu and Binoop Nanu for early prototyping. We acknowledge the EBI cloud consultants Santiago Insua and David Ernesto Gómez Gutiérrez for their help in deploying the website. We are grateful to the IMPC production working group for their work that made it possible to display the ES Cell and CRISPR alleles on the genome browser. We would also like to acknowledge NIH programme directors Colin Fletcher and Oleg Mirochnitchenko and programme analysts Maya Vanzanten and Sofia Martin for their help and support.

## Funding

National Institutes of Health (NIH) [2UM1HG006370-11, 5UM1HG006370-13, 3UM1HG006370-12S2, 1U24OD038424-01]; EMBL-EBI Core Funding. Funding for open access charge: NIH.

## Conflict of Interest

None declared.

## Notes

### Competing Interest Statement

The authors have declared no competing interest.

